# Domain-combination based pan-genomic Tree of Life

**DOI:** 10.1101/2025.10.16.682753

**Authors:** Rubén García-Domínguez, Carlos Santana-Molina, Damien P. Devos

## Abstract

Reconstructing the Tree of Life (ToL) remains a central challenge in evolutionary biology. While sequence-based methods have resolved much of life’s diversity, deep evolutionary relationships often remain elusive, particularly across the three superkingdoms of life. Here, we present a complementary approach using genome-wide occurrence of protein domain combinations as phylogenetic signal. We analyzed 5,343 complete proteomes spanning Bacteria, Archaea, and Eukaryota, and inferred a pan-genomic ToL from ∼77,000 domain combinations, including isolated domains and domain pairs. Our reconstruction recovers major clades and captures both vertical inheritance and signals from non-vertical processes such as lateral gene transfer, endosymbiosis, and genome reduction. Ancestral reconstructions reveal functional innovations that mark key evolutionary transitions—such as the emergence of vesicle trafficking in Eukaryota or methanogenesis in Archaea. We found that organisms with reduced genomes, like Chlamydiota, tend to be misplaced in domain combination-based phylogenies due to sparse combinatorial data. Our results demonstrate that domain combinations carry a strong, evolutionary signal that is both biologically and functionally informative. While not a substitute for sequence-based phylogenetics, domain combination-based trees offer an alternative perspective, capable of integrating both vertical and lateral processes into a unified evolutionary framework.

**Author Summary:** Traditional evolutionary trees are typically built from sequences, but this approach struggles to resolve deep branches in the Tree of Life. Here, we use combinations of protein domains—the functional modules that make up proteins to resolve major relationships in the tree. By analyzing over 5,000 complete proteomes from Bacteria, Archaea, and Eukaryotes, we reconstruct a Tree of Life that reflects both vertical inheritance and key evolutionary processes like endosymbiosis and gene transfer. This domain-based approach not only recovers known relationships but also reveals how major biological innovations emerged. Our results show that domain combinations provide a powerful and interpretable signal for studying evolution at a genomic scale.

## Introduction

The tree of life (ToL) is the main organizing principle of biology [1, 2]. Its reconstruction remains one of the most studied—and most disputed—questions in evolutionary biology. While sequence-based methods have resolved much of life’s diversity, deep relationships remain unresolved. Traditional phylogenetic methods have been instrumental in reconstructing evolutionary relationships, but they face methodological hurdles, particularly for deep phylogenies and when evolutionary rates vary across sequences (heterotachy) [3], when aligning highly divergent sequences, or when relying on a limited set of universally conserved marker genes. The necessity of universal conservation often restricts large-scale analyses to a handful of proteins, capturing only a fraction of genomic content—a limitation sometimes referred to as the ‘tree of one percent’ [4]. Additionally, the pervasive effects of horizontal gene transfer (HGT) and signal saturation can introduce inconsistencies that complicate phylogenetic reconstruction. Here we propose that protein-domain combinations offer a complementary approach that is less affected by these limitations.

Whole-genome and whole-proteome phylogenies have emerged as complementary approaches that extend beyond traditional sequence-based methods [5–7]. By leveraging broader genomic features—including ortholog content, k-mers, and protein domains—these approaches provide alternative perspectives on evolutionary relationships. Domain-based phylogenies, in particular, integrate structural and functional signals, offering a higher signal-to-noise ratio and reducing the need for manual intervention steps such as orthology assignment and alignment trimming [8].

Protein domains are the fundamental building blocks of proteins, defining their structure, function, and evolutionary history. The composition and arrangement of domains largely determine protein function, and the emergence of new domain architectures is a key driver of protein evolution [9, 10]. Studies suggest that the gain of novel domain architectures differentiates eukaryotes from prokaryotes, marking evolutionary transitions such as the origin of multicellularity [7, 11–13]. Despite a near-infinite theoretical space of combinations, only a limited subset is observed across genomes—implying that natural selection shapes domain architectures [7–9, 14]. In a linguistic metaphor, domain architectures have been described as the “words” and “sentences” of a genomic language, where each new arrangement expands an organism’s functional repertoire [7]. Given their status as discrete evolutionary units, domains and their combinations provide a robust phylogenetic framework. Beyond their structural and functional informativeness, domain architectures offer an orthogonal lens on vertical inheritance, horizontal transfer, and convergence—complementing traditional sequence-based signals and allowing for a broader exploration of evolutionary patterns.

Here, we present a pan-genomic Tree of Life built from 5,343 proteomes spanning Bacteria, Archaea, and Eukaryota. Using Pfam domains and domain pairs as binary phylogenetic characters, our reconstruction not only recovers established clades but simultaneously reveals lineage-specific functional innovations. Unlike classical trees, our domain-combination approach highlights patterns of endosymbiosis, horizontal gene transfer (HGT), and vertical inheritance—offering a broader, complementary perspective on evolutionary history. We also reconstruct ancestral domain combinations, revealing how specific innovations map onto key evolutionary transitions across the tree. Together, these results support domain-based phylogenies as a robust and interpretable alternative framework—complementary to sequence-based methods—for reconstructing evolutionary relationships and functional diversification.

## Results and Discussion

### 1. Proteome and Functional Domain Features Across the Tree of Life

We analyzed 5,343 proteomes within the three superkingdoms of life (Fig 1A), including 3,149 Bacterial (78 phyla), 1,367 Archaeal (16 phyla), and 827 Eukaryotic proteomes (21 phyla), to explore patterns of proteome diversity and functional domain architecture. Despite the lower number of eukaryotic representatives, their proteomes dominate in terms of amino acids counts over Bacteria and Archaea (Fig 1.A3) and appear as intermediate in terms of protein counts (Fig 1.A2). Our dataset covers a broad range of organisms, encompassing diverse ecological and evolutionary contexts, although it reflects a known bias favoring more extensively studied lineages. We analyzed multiple metrics (Fig 1B–G), including completeness, total proteins, domain diversity, domain-pair diversity, and combinatory rate (defined as pairs diversity divided by domain diversity).

**Fig 1.**
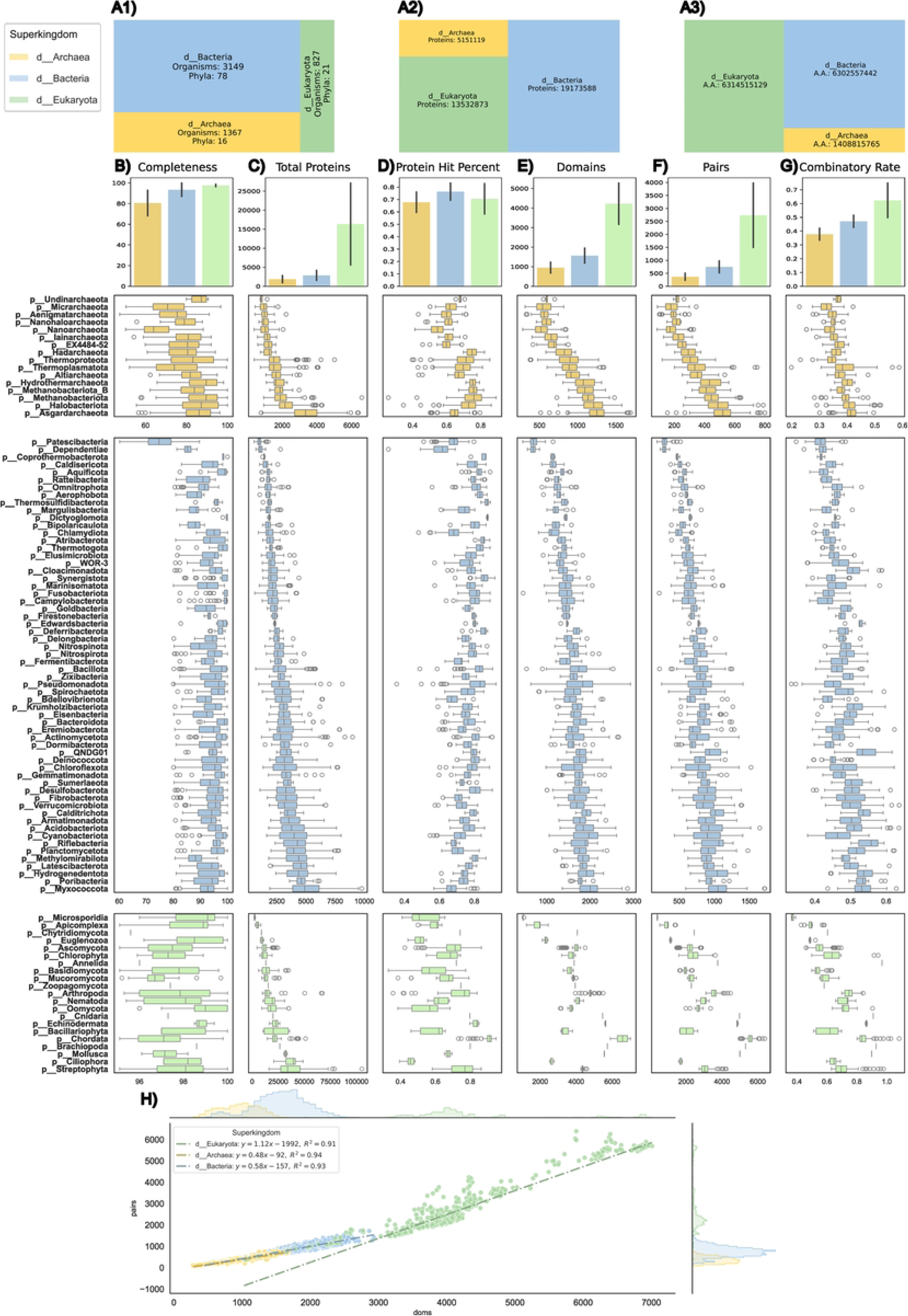
Proteome-wide summary and comparative distributions. A)Proteome overview. Tree-map showing the total number of proteomes (A1), protein (A2) and amino acids (A3), broken down by superkingdom (Eukaryota, Archaea, Bacteria) and colored accordingly; B–G. Quantitative feature distributions per phyla. Boxplots in panels B–G are grouped by superkingdom (gold = Archaea; blue = Bacteria; green = Eukaryota): B) Genome completeness (%) as estimated by BUSCO and GTDB for Eukaryotes and Prokaryotes (Bacteria and Archaea) respectively. C) Total number of predicted proteins. D) Fraction of proteins with at least one Pfam domain hit. E) Total number of unique Pfam domains. F) Total number of unique domain-pairs. G) “Combinatory rate”—the ratio of unique domain-pairs to unique domains; H. Domain–pair versus domain count regression. Scatterplot of unique domains (X) against unique domain-pairs (Y) for each proteome, colored by domain of life, with marginal histograms. Linear fits and corresponding equations (slope ± SE, R²) are overlaid for each superkingdom.

Our results reveal patterns in genome completeness among the three superkingdom, although these patterns are influenced by the criteria used for dataset selection (Fig 1B, see Methodology). In terms of proteome size, Eukaryotes exhibit the largest protein counts, consistent with their increased genome complexity and functional capacity (Fig 1C). In contrast, Bacteria and Archaea display smaller proteomes, often influenced by genome reduction in specific lineages, such as Patescibacteria, Dependientae and Chlamydiota in Bacteria, and many DPANN phyla like Undinarchaeota and Micrarchaeota in Archaea (Fig 1C).

The Protein Hit Percent (PHP) (Fig 1D), the percentage of proteins assigned with at least one Pfam domain, is highly variable across all superkingdoms. In Eukaryotes, it likely reflects variable proportions of disordered or non-globular regions and a bias toward model organisms [15]. In Bacteria and Archaea, PHP variability is narrower yet notable, with reduced-genome lineages such as Patescibacteria and several DPANN archaeal phyla displaying notably lower PHP.

Functional diversity, as measured by Pfam domains and domain pairs, was highest in Eukaryota, highlighting their unique ability to innovate functional repertoires (Fig 1E,F). This is particularly notable in multicellular eukaryotes, such as Chordata and Arthropoda, which display high levels of domain pairs compared to Bacteria and Archaea (Fig 1E,F). Among Bacteria, phyla like Myxococcota, Hydrogenedentota or Riflebacteria show elevated levels of domain and domain pairs, while reduced-genome lineages like Patescibacteria, Chlamydiota, and Dependientae exhibit significantly fewer unique domains and combinations (Fig 1E,F). Archaea displayed lower diversity of domains and pairs, with Asgardarchaeota and Halobacteriota forming an intermediate group, overlapping with the lower range of bacterial diversity but still lagging behind most Bacteria (particularly those without significant genome reduction) and Eukaryotes in pairs innovation innovation (Fig 1E,F).

The relationship between the number of domain pairs and total domains, defined as the Combinatory Rate (Fig 1G), highlights clear evolutionary trends. Archaea show a nearly flat regression slope, indicating minimal domain combination, while Bacteria exhibit a steeper slope, reflecting greater domain pairing (Fig 1H). Eukaryotes display the most pronounced slope (Fig 1H), underscoring their ability to innovate through extensive domain combinations. This is particularly evident in multicellular Eukaryotes, such as Metazoa and Streptophyta (S1 Fig), which exhibit exceptionally high rates, correlating with their increased functional complexity. These results reinforce the idea that domain combination is a major driver of functional innovation and evolutionary adaptability across the three superkingdoms of life, particularly in Eukaryotes [9, 12, 16].

### 2. Unprecedented taxonomic resolution by pan-domain Tree of life

We constructed a pan-domain Tree of Life using a novel approach based on minimal combinations of protein domains, including isolated domains and domain pairs (Uni-domain content + pair content). In order to estimate the best use of our dataset, we compared it against other strategies (Fig 2A), such as trees based solely on domain content or combined domain and pair content. Those trees were inferred from occurrence matrices of domain combinations, employing binary character substitution models (GTR2) as implemented in IQ-TREE. The domain content tree (S2 Fig) showed weaker overall relationships among major supergroups, in agreement with previous analyses [8, 17]. However, this dataset yielded more coherent placement of reduced-genome lineages like Chlamydiota, and Dependientae, among others, likely due to the higher proportion of single-domain proteins in these lineages as well as the influence of a lower sparsity of the matrix. On the other hand, combining domain content and pair content increased redundancy and computational cost without significant improvement in resolution (S3 Fig). Thus, we focused on domain combinations (Uni-domain + pair content) as the optimal solution [14]. Our domain-combination tree is based on ∼18M proteins, from which we defined ∼77K combinations (15.9K isolated domains + 61.1K pairs of domains), considering combinations that appear at least in 2 organisms, to avoid singletons (combinations only present in one organism). Those numbers highlight the importance of architecture (represented by pairs of domains) in evolutionary reconstructions.

**Fig 2.**
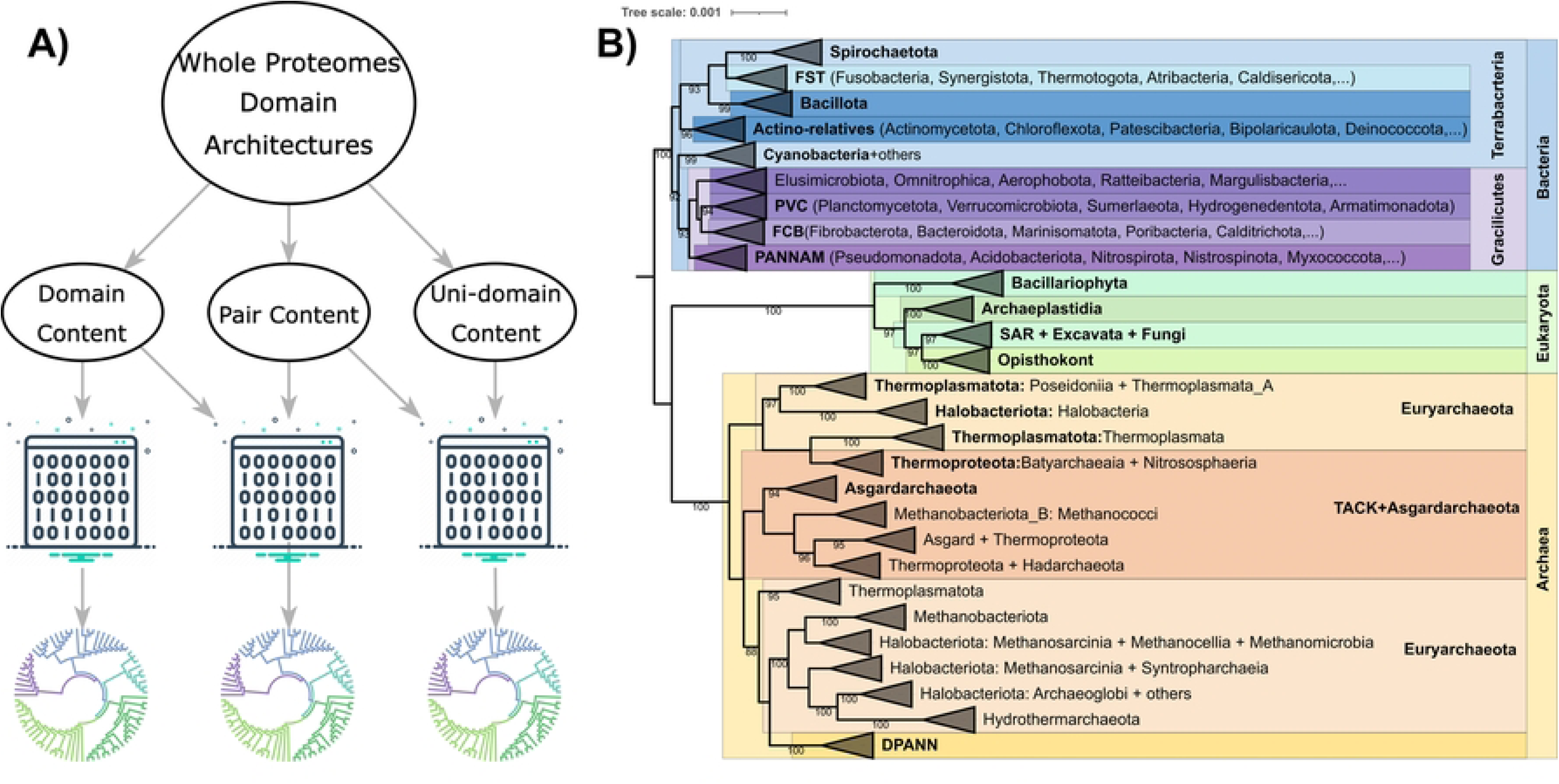
Construction and final result of the combination-based Tree of Life. A) Scheme of the different approaches used to create the different binary matrices based on different features, which were used to elucidate domain-based phylogenies. B) Collapsed ML-Tree done using IQ-TREE with binary substitution model (GTR2+F0+R10), ultrafast bootstrap 1,000 iterations and –bnni option (UFBoot trees by nearest neighbor interchange (NNI)) using combined pair content + uni-domain content binary matrices as input. Different superkingdoms appear with different colors; Blue, Bacteria; Green, Eukaryota; Yellow, Archaea. Supergroups and different clades within each superkingdom display color variations of the superkingdom color.

The resulting tree aligns well with the classical 3-kingdoms topology of the Tree of Life (Fig 2B; [2, 18–20], recovering Bacteria, Archaea, and Eukarya as distinct monophyletic groups. Within Bacteria, the clear separation of Gracilicutes and Terrabacteria reflects established phylogenetic relationships, with superphylum like PANNAM [21] (Pseudomonadota, Acidobacteriota, Nitrospirota, Nitrospinota, Aquificota and Methylomirabilota), PVC, and FCB robustly resolved among Gracilicutes and FST and Actinomycetota-relatives clustering confidently among Terrabacteria [21–23]. Archaeal lineages, including DPANN, Crenarchaeota, and Euryarchaeota, also match traditional topologies while capturing patterns of genome reduction and niche adaptation. Besides, our reconstruction accurately matches the recent position of DPANN diverging from within Euryarchaeota [24], instead of being positioned as a basal group to the whole Archaea, highlighting that the method is less affected by strong divergence [25]. In Eukaryota, major clades like Opisthokonta, Archaeplastida, and SAR are strongly recovered while showing some discrepancies with groups like Excavata appearing within SAR, nevertheless, underscoring the method’s capacity to resolve complex evolutionary relationships.

These results validate our pan-domain tree as a robust alternative for tracing evolutionary relationships and functional dynamics across the three superkingdoms. However, the method is not without inconsistencies, raising important questions about specific phylogenetic placements and functional overlaps.

### 3. Pan-domain Tree of Life illustrates evolutionary relationships beyond vertical evolution

Although vertical inheritance typically shapes evolutionary histories, processes such as lateral gene transfer (LGT), endosymbiosis, and genome reduction introduce confounding evolutionary signals [26–29]. Rather than mere methodological biases, these signals can reveal biologically significant events shaping lineage evolution. Here, we highlight three examples illustrating how explicitly addressing these non-vertical processes provides insights into key evolutionary adaptations and relationships: (1) intra-archaeal topology influenced by LGT acquisitions from bacteria, (2) archaeal-eukaryotic relationships shaped by endosymbiosis, and (3) reduced-genome lineages like Chlamydiota affected by genome reduction which therefore result in increased data sparsity. In all cases, we applied a Weighted Site Likelihood (WSL) approach to compare alternative topologies and to determine the supporting domain combinations [30].

#### LGT acquisition from Bacteria affects archaeal topology

Halophilic archaeal groups, such as Halobacteria (within Halobacteriota) and Poseiidonia (within Thermoplasmatota) as well as some classes of the phylum Thermoproteota, appear paraphyletic and close to the archaeal root in our pan-genomic tree (Fig 3A Left). To evaluate the domain combinations supporting each topology, we compared a topology relocating these groups to Euryarchaeota and TACK + Asgardarchaeota respectively (Fig 3A Right). Weighted site likelihood (WSL) analyses reveal that the domain-based topology is supported by domain combinations linked to potassium channels, osmoregulatory proteins, and other potential niche-specific adaptations related proteins (Fig 3B). The respective phylogenies of some of these proteins, reflect LGT events from bacteria to these specific archaeal clades explaining the sharing of domain between these, resulting in the attraction of these archeal lineages to the base of archaea (Fig 3C1). In contrast, the topology manually modified to reflect a more canonical archaeal clades distribution is associated with domain combinations related to DNA-binding and core cellular functions, suggesting that these functions support vertical inheritance relationships (Fig 3B, 3C-2). Sequence-based phylogenies constructed from domain sets favoring each topology reveal contrasting evolutionary dynamics: those supporting the domain-based arrangement, inferred from Chlorite_dismutase (PF06778) and COX15-CtaA (PF02628), exhibit extensive LGT topology, while those favoring the canonical topology, based on Ribosomal_S4 (PF00163) and the Topoisom_I_N–Topoisom_I pair (PF002919-PF01028), align with core archaeal traits and a more conserved vertical heritage. Those results highlight the role of lateral gene transfer (LGT) in shaping archaeal domain-based phylogenies (Fig 3C-1) in agreement with previous analyses [31–33].

**Figure 3.**
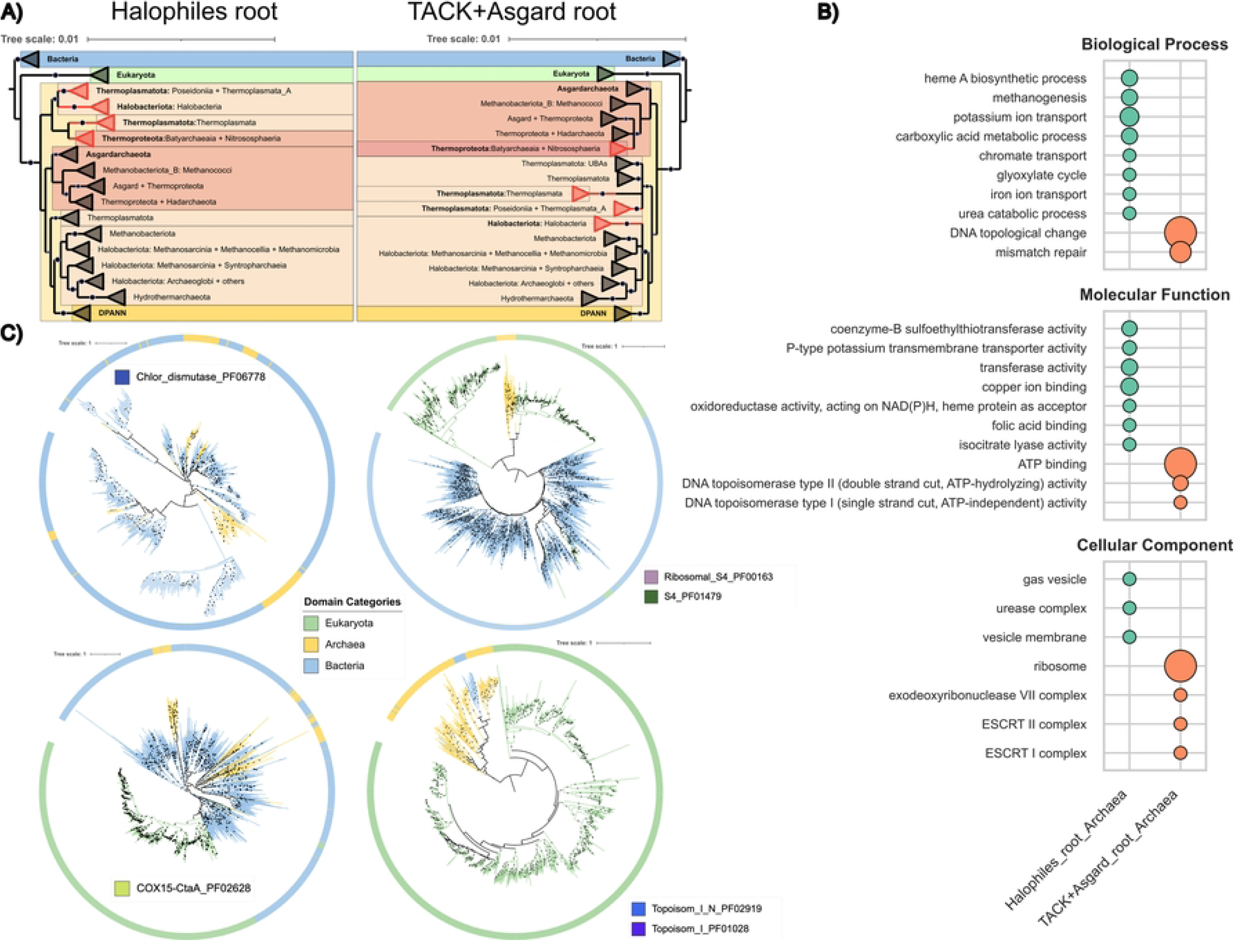
WSL analysis of archaeal topology. A) Original topology of the domain combination based tree with the “missplaced” clades in red and the manually modified topology of the tree. Both were used to calculate the ΔWSL and select the combination that most favored each topology. C) Sequence-based phylogenies of selected domain combinations that favour each topology. B) Functional enrichment analysis of each cluster of domain combinations.

#### Endosymbiosis: Archaea-Eukaryota Relationship

The domain-combination-based phylogeny recovered Eukaryota as a sister group to Archaea [34], differing from the widely accepted view that Eukaryota emerged from within the Asgard clade (Fig 4A, left and right, respectively). This discrepancy likely stems from differences in the sizes of their domain combination repertoires—Eukaryota exhibiting significant expansion and Archaea showing notable reduction—as well as potential methodological biases introduced by including Bacteria. When Bacteria are removed, Eukaryota and Asgardarchaeota appear closer, which could even be interpreted as Eukaryota within Asgardarchaeota, depending on the root position (S4 Fig), this suggests that bacterial lineages introduce confounding signals/phylogenetic attractions when resolving deep archaeal-eukaryotic relationships. Using a Weighted Site Likelihood (WSL) approach to compare alternative topologies (Fig 4B), we find that domain combinations supporting the Eukaryota-Archaea sisterhood are enriched for mitochondrial and chloroplastic proteins, highlighting the influence of endosymbiosis on the topology of our pan-genomic Tree of Life. Conversely, domain sets favoring the Eukaryota-within-Asgard topology show weaker enrichment for these endosymbiotic markers, indicating a stronger vertical inheritance signal. These findings underscore the role of endosymbiotic-driven adaptations in shaping phylogenetic trees and emphasize the need to account for these influences when interpreting deep evolutionary relationships.

**Fig 4.**
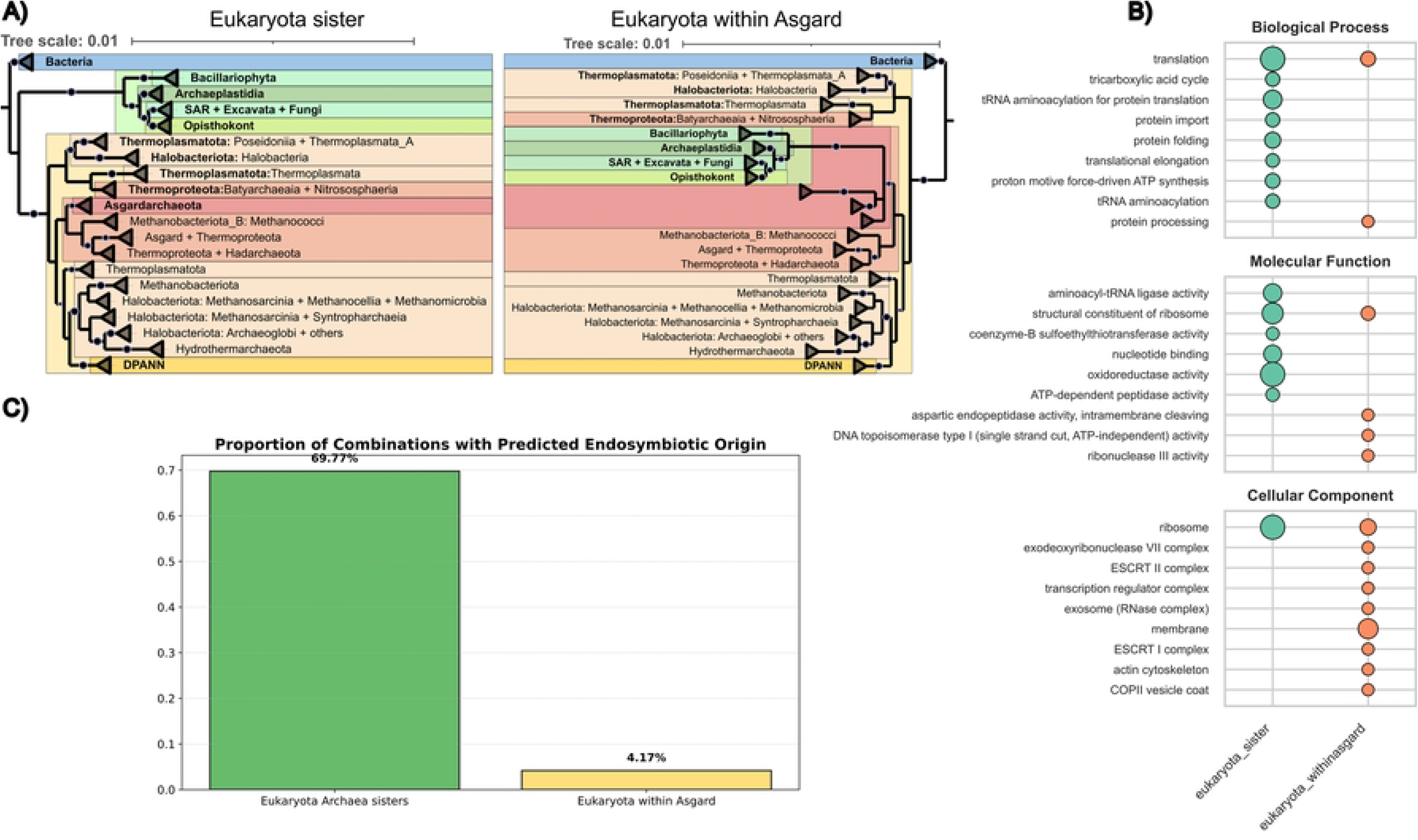
WSL Analysis of Archaea-Eukaryota topology. A) Original topology of the domain combination based tree and the manually modified topology of the tree with the whole eukaryotic clade placed within Asgardarchaeota. Both were used to measure the WSL and calculate the ΔWSL and select the combination that favored the most each topology. hB) Percentage of domain combinations from each subgroup that have more than 50% of their protein sequences predicted to be chloroplastic or mitochondrial using targetP. C) Functional enrichment analysis of each cluster of domain combinations.

#### Reduced Genome and sparseness issue: Chlamydiota

Chlamydiota, traditionally classified within the PVC superphylum, appears instead within the PANNAM cluster in our domain-combination-based tree, likely due to its reductive evolutionary trajectory and the effects of long-branch attraction (LBA). This misplacement improves when using a domain-content tree (S2 Fig), where the matrix size is significantly reduced (∼14K in the domain-content tree vs. ∼77K in the domain-combination tree). The increased sparsity in the combination matrix disproportionately affects reduced proteomes, as they accumulate large numbers of absences, potentially distorting their phylogenetic placement. WSL analysis shows that the PANNAM placement (Fig 5A) is supported by domain combinations enriched in conjugative elements, such as TraN and TraE, suggesting historical gene exchange and functional convergence with other PANNAM members [35]. However, the topology placing

**Fig 5.**
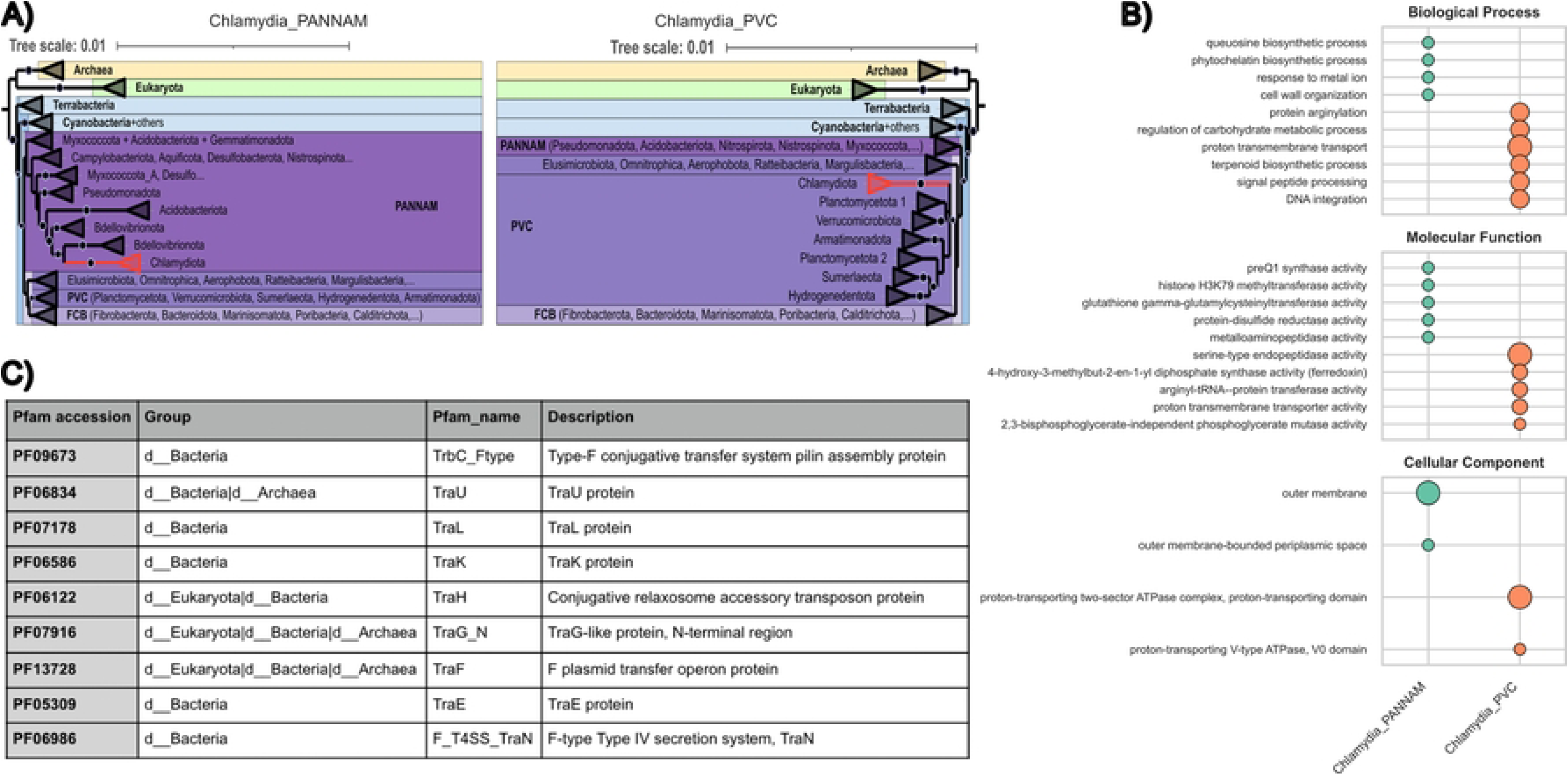
WSL Analysis of Chlamydiota placement. A) Original topology of the domain combination based tree with the “missplaced” clades in red (Chlamydiota, within PANNAM) and the manually modified topology of the tree (Chlamydiota within PVC superphylum). Both were used to calculate the ΔWSL and select the combination that favored the most each topology. B) Table of domain combinations (in this case all isolated domains) with Tra-annotations, related with conjugation plasmids. C) Functional enrichment analysis of each cluster of domain combinations.

Chlamydiota within PVC (Fig 5A) is supported by domain combinations reflecting core metabolic and cellular machinery, with WSL analysis favoring domain combinations linked to conserved biological processes, including ATPase complexes and terpenoid biosynthesis. This contrast highlights the challenge of resolving relationships for lineages with important genome size differences, where both adaptive processes (e.g., LGT and functional convergence) and vertical inheritance influence evolutionary patterns.

### 4. Ancestral reconstruction of domain combinations reveals lineage-specific functional innovations

The topology of our domain-combination based ToL further allowed us to explore how protein-domain combinations have diversified across the Tree of Life, we reconstructed ancestral combinations on our phylogeny via maximum-likelihood and annotated each combination with Pfam2GO. We then quantified, at every internal node, the total number of combinations inherited from its parent versus those unique to that lineage (grey bars for root-derived, intermediate colours for deeper ancestors, top segment for node-specific innovations) (Fig 6B). The results revealed that LUCA already harboured ∼2,800 domain combinations, but the eukaryotic stem contributed the largest—over 1,500 novel architectures—followed by Bacteria (∼500) and more modest gains in Arkarya and Archaea (∼300 and ∼142 respectively). This indicates that the LUCA was already complex, that Eukaryota additionally developed roughly only half of the previously existing architecture repertoire and that Archaea are the least inventive, or have evolved by many losses, as suggested previously [29].

**Fig 6.**
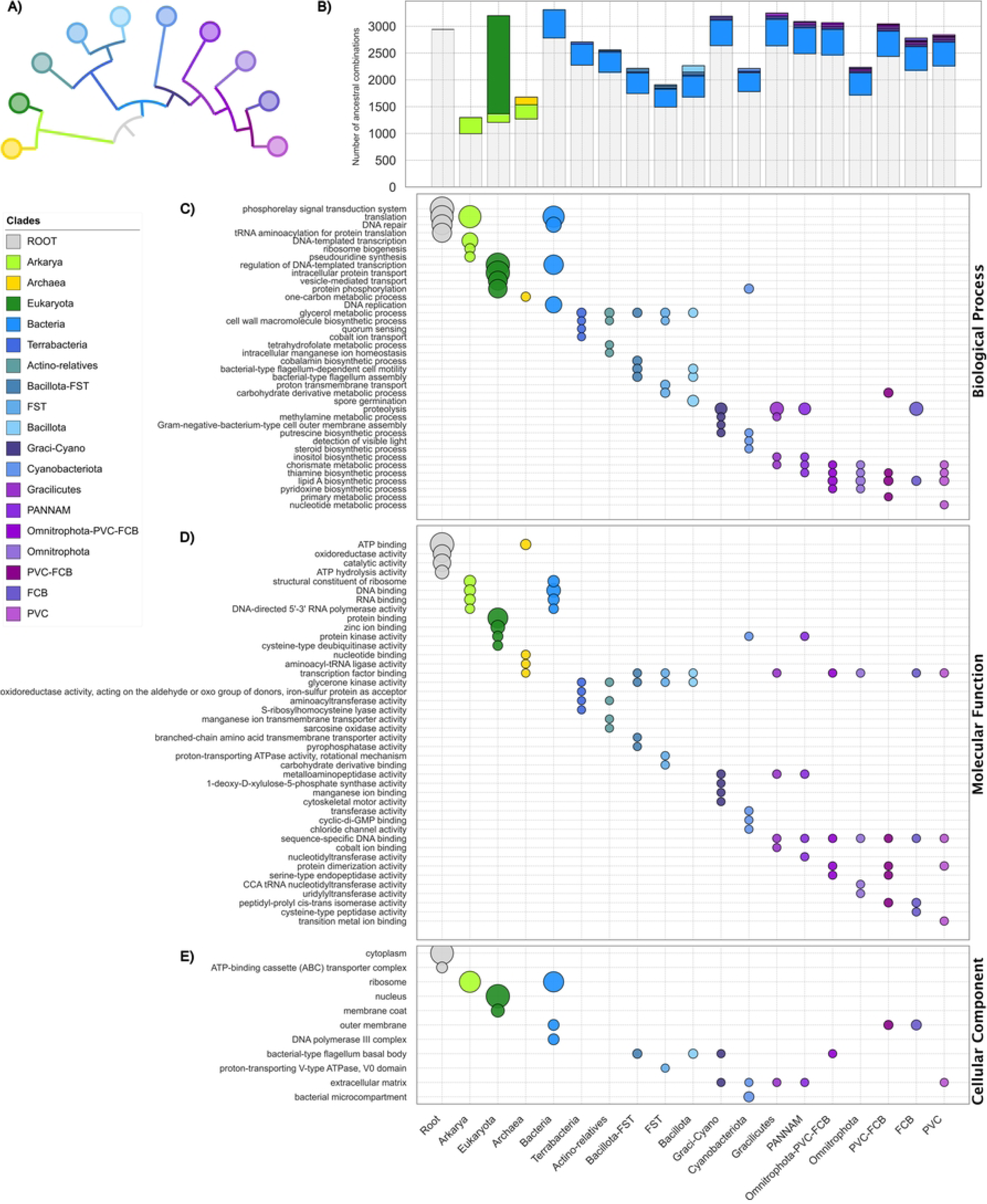
Ancestral reconstruction of protein-domain combinations and functional enrichment. (A) Phylogenetic tree of major clades, colored by lineage, including ancestral nodes. (B) Bar-plot showing, for each ancestral node (both internal and tips), the total number of inferred domain combinations from the root (grey), those inherited stepwise from each intermediate ancestor (coloured segments, in phylogenetic order), and finally the combinations unique to that node (top-segment, matching the node’s colour). (C–E) GO-term enrichment for each ancestral domain-combination set, split by aspect. For each GO aspect, we display the four most significant terms, bubble sizes according to their count in the node’s set of functions: (c) Biological Process, (d) Molecular Function, (e) Cellular Component. For each node we show the four most significant GO terms (lowest p-values), with bubble size proportional to the count of that GO term in the domain set and bubble colour indicating the node’s clade.

We note that, due to bacterial genomes being overrepresented in our dataset (Fig 1A), our ancestral reconstructions may introduce some biases toward bacterial ancestrality, which could account for the higher prevalence and retention of root-derived traits in Bacteria compared to Arkarya and its descendants. Alternatively, this pattern might also reflect that modern bacterial domain combinations are, on average, more similar to the ancestral state than those of archaeal lineages, preserving a greater share of ancient combinations.

Next, we tested for functional enrichment in the “novel” ancestral domain-combinations at each node by comparing them to all ancestral combinations within the same subtree, using one-tailed Fisher’s exact tests on GO annotations (Fig 6C–E).

The Last Universal Common Ancestor (LUCA) was enriched in oxidoreductase functions, highlighting the role of redox reactions in early metabolism, and general catalytic folds (e.g., NADH_dehydrogenase_N PF00146; MMR_HSR1 PF05188; cytochrome b C-terminal PF00032; P450 monooxygenase PF00067). It also carried basic protein phosphorylation/kinase modules, pointing to ancient redox chains and simple regulatory circuits. Moreover, LUCA possessed ancestral phosphorelay transduction capabilities—early sensor modules such as PAS domains (PF00989) and GAF domains mediating phosphate transfer. We also observe more specialized two-component modules that became fixed in Bacteria (e.g., Response_reg|HATPase_c fusions like PF00072–PF02518). Finally, LUCA’s toolkit included ABC-type transporters (ATP-binding cassette complexes), underscoring an early need for active import–export across its membrane.

The Arkarya ancestor boosted translation and RNA processing—ribosomal modules Ribosomal_S13_N–S15 (PF08069–PF00312), RNA_pol_Rpb2_3–Rpb2_4 (PF04565–PF04566), RNA-binding KH_1 (PF00013) and tRNA-modification THUMP (PF02926)—and enriched structural constituents of the ribosome and ribosome, underscoring an early shift toward a more fine-tuneable protein-synthesis machinery that would underpin complex gene expression in Archaea and Eukaryota, in agreement with the know closer relationship between archaea and eukaryotes on their information manipulation machineries.

Within Archaea, core ribosomal and tRNA-aminoacylation domains persisted—such as the Thr-specific ligase module (PF08915–PF00587)—while new transcription-factor binding modules and one-carbon metabolic enzymes—such as the MtrH methyltransferase subunit (PF02007) and the MCH cyclohydrolase (PF02289)—linked to archaeal methanogenesis via the tetrahydromethanopterin pathway, driving key methyl-group transfers that power methane production under anaerobic conditions [36, 37], in agreement with this strictly archaea process.

The eukaryotic ancestor unveiled vesicle-trafficking innovations (Clathrin_lg_ch PF01086; Sec1 PF00995; Adaptin_N|Alpha_adaptinC2 PF01602–PF02883; Coatomer_WDAD|COPI_C PF04053–PF06957), elaborated membrane-coat complexes and nuclear-pore architectures (NUC153 PF08159; Nuclear pore anchor NAP PF00956; Mpp10 PF04006), and incorporated diverse protein-kinase activities (Protein kinase PF00069; Protein tyrosine kinase PF07714; Ser/Thr kinase active site PF00509), highlighting the importance of membrane-based organelles in eukaryotes and their complex signaling pathways.

Bacteria exhibit a strong signal for DNA/RNA functions (RNA_pol_Rpb2_2–Rpb2_3 PF04561–PF04565; Phage_integrase PF00589; Bac_DNA_binding PF00216) and conserved FtsK-type partitioning (PF01580–PF09397; PF17854–PF01580). They also carried outer-membrane biogenesis domains such as POTRA–Omp85 (PF07244–PF01103), in agreement with the proposal that early Bacteria were diderm, not monoderm [38].

Terrabacteria were marked by glycerol metabolism and cell-wall synthesis domains (FemAB–FemAB PF02388–PF02388), reflecting the reliance of monoderm Firmicutes on the femAB operon for pentaglycine interpeptide bridge formation in peptidoglycan—a key determinant of cell-wall rigidity and methicillin resistance [39].

Gracilicutes were characterized by an expansion of extracellular metallopeptidases (Matrixin Peptidase_M10 PF00413; cytosolic aminopeptidase Peptidase_M17 PF00883) and specialized methylamine-utilization enzymes (MauE PF07291), alongside enrichment of membrane-assembly factors (LptE PF04390) and flagellar switch components (FliM|FliN PF02154–PF01052). Together, these innovations underpin their ability to remodel the cell surface, exploit simple nitrogen sources, build complex outer envelopes and power motility.

Within Bacillota, spore-germination modules such as the GerA sensor (PF03323), Spo0A receiver–DNA-binding fusions (PF00072–PF08769) and dedicated germination proteases (PF03418) mark the advent of cellular dormancy and stress resilience. Cyanobacteriota are defined by elaborate microcompartment assemblies (BMC PF00936; EutN_CcmL PF03319).

Finally, the Omnitrophota–PVC–FCB cluster shows LpxC–FabA fusions (PF03331–PF04977) at the nexus of lipid A and phospholipid synthesis—potentially reflecting a primitive coupling of these two pathways. Even when unfused, LpxC and FabZ compete for the common precursor R-3-hydroxymyristoyl-ACP, and LpxC levels are controlled by FtsH/YciM-mediated proteolysis to balance lipopolysaccharide and phospholipid production [40, 41].

## Conclusions

Our domain-combination-based Tree of Life demonstrates that protein domain combinations capture a strong vertical evolutionary signal across the tree of life. Despite relying on presence-absence patterns rather than sequence alignments, the reconstructed phylogeny is remarkably robust and broadly consistent with established taxonomies. This highlights the potential of domain combinations as effective phylogenetic markers, preserving core evolutionary relationships even over deep evolutionary timescales.

Importantly, our approach also captures signals beyond strict vertical inheritance, including signatures of lateral gene transfer, endosymbiosis, and genome reduction. Rather than being viewed as methodological artifacts, these patterns reflect biologically meaningful processes that have shaped organismal evolution and are well documented in the literature. Thus, domain-based phylogenies offer a unique opportunity to integrate both vertical and non-vertical processes into a comprehensive evolutionary framework.

Nevertheless, the method has limitations, particularly regarding reduced genomes, where sparsity can distort phylogenetic placement. Future work will focus on addressing these challenges by extending the framework to structural domains, which typically offer broader proteome coverage, minimize annotation biases, and further reduce the need for manual curation steps.

Finally, while domain combination phylogenies bypass some of the bottlenecks faced by classical sequence-based approaches—such as orthology assignment, alignment ambiguity, and marker selection—they should not be seen as a replacement. Instead, they represent a complementary perspective, capable of capturing both the canonical tree-like evolution and the complex reticulate processes that shape life’s history. This integrated view opens new approaches to detect, interpret, and understand major evolutionary innovations across the Tree of Life.

## Materials and Methods

### 1.- Dataset selection

For this study, a total of 5,343 proteomes were obtained from different sources. Prokaryotes (3,149 Bacteria and 1,367 Archaea) were obtained from GTDB [42] by the selection of representative proteomes. For bacteria, we sampled proteomes at order level: if the number of orders within a phylum is higher than 100, we selected one proteome per order with the higher completeness, the lower contamination, and type-strain if possible (those with genome completeness lower than 80% and contamination higher than 5% were excluded, with the exception of Patescibacteria for which we included genomes with completeness higher than 60%). For those phyla in which the number of orders is lower than 100, we sampled at genus level, by selecting the top quality genomes with the same threshold as above. If the sampled genomes for a phylum was still lower than 50 we made a second round of sampling. The rationale of this iterative sampling was to achieve a taxonomically balanced and complete dataset. For archaea, due to genome reduction being more prominent specially in DPANN organisms, we selected genomes by sequence identity of the concatenated gene markers provided by GTDB and which exclusively contains representative taxa. We clustered the concatenation by taxonomic classes, and reduced redundancy of the concatenation at sequence identity of 85% using trimAL (-maxidentity 0.85), using the remaining proteomes as representative for our final dataset. Eukaryotes (827) were obtained from Uniprot Reference Proteomes, considering a stringent filter of 95% of BUSCO completeness score [43].

### 2. Dataset Preparation

To identify protein domains within each proteome, the pfam_scan.pl script from the Pfam database (http://ftp.ebi.ac.uk/pub/databases/Pfam/Tools/ version 2017-03-01) was employed. Pfam provides curated multiple sequence alignments for protein families, leveraging hidden Markov model profiles to detect domains in new sequences [15]. Only domains, families and repeat with an E-value < 0.001 were considered significant, following established methodology in the field [7].

The output from pfam_scan.pl was processed to determine the domain architecture for each protein. In case of overlapping domains (>30% overlap), the domain with the highest bitscore was retained, ensuring the most biologically relevant annotation [7]. This approach minimizes redundancy in domain assignment while maintaining accuracy, allowing for the robust characterization of protein domain architectures across proteomes.

### 3. Quantitative Analysis

Several quantitative metrics were computed to characterize the diversity and functional capacity of organisms across different clades. These metrics provided a foundation for evaluating evolutionary patterns and functional innovations across the dataset, as illustrated in Fig 1.

● **Domain diversity**: The total number of unique domains in each proteome was quantified.
● **Domain pairs**: The number of domain pairs within each proteome was calculated to assess combinatorial diversity, taking into account the order of the domains (i.e. pairs AB and BA are counted as two different pairs).
● **Genome completeness**: Completeness scores, domain coverage, and other structural features were analyzed.

### 4. Binary Matrices for Phylogenetic Reconstruction

We generated multiple binary matrices to construct the pan-domain Tree of Life, each representing different types of information:

- **Domain content matrix**: A binary matrix indicating the presence or absence of individual domains across proteomes. This approach works well for reduced-genome lineages as it minimizes matrix sparsity, which otherwise disproportionately impacts organisms with highly reduced genomes. Such organisms, already characterized by a high proportion of missing data (many zeros), are more affected by sparsity, leading to issues like Long Branch Attraction (LBA). This can result in poorer phylogenetic placement.
- **Domain pair content matrix:** It captures domain combinations, offering a richer view of proteome content. However, this approach increases sparsity and redundancy without significantly improving phylogenetic accuracy. The increased number of alignment columns also leads to longer computational times.
- **Combined matrix:** It integrates domain pair content with uni-domain architectures (proteins containing only one domain). This approach maximizes proteome coverage while mitigating the increase in sparsity, providing better global placement for non-reduced genomes and improving the resolution of taxonomic groups.

The matrices were used as input for phylogenetic reconstruction with IQ-TREE [44], employing binary substitution model GTR2+F0+R10, selected via ModelFinder [45] and ultrafast bootstrap approximation (UFBoot2) [46]. In this model:

- GTR2 is a general time-reversible binary substitution model designed for presence/absence data.
- F0 specifies the use of empirical state frequencies.
- R10 represents the number of rate categories in the FreeRate model, allowing for site-specific rate variation.

The –bnni option was used to optimize branch lengths and topologies, improving convergence and mitigating model violations. The ultrafast bootstrap method (-bb 1000) was employed to estimate branch support, offering computational efficiency and reliable confidence intervals in large datasets.

### 5. Weighted Site Likelihood (WSL) Analysis

To compare topologies derived from our domain-based approach with alternative phylogenies (both classical and manually curated), we utilized a Weighted Site Likelihood (WSL) approach [30]. This method allows for the identification of domain combinations with the highest likelihood contributions to each topology, enabling direct comparisons between topological models. WSL is particularly useful for interpreting the biological relevance of specific domain combinations, as demonstrated in earlier studies [30]. By extracting functional combinations linked to specific topologies, WSL provides insights into the evolutionary forces shaping domain architectures.

### 6. Functional Annotation with Pfam2GO

To avoid biases introduced by protein-based functional annotations, we leveraged Pfam2GO to assign functional annotations directly to domain combinations. This method ensures functional interpretation is strictly domain-centric, avoiding biases from full-length protein annotations. A custom pipeline was developed to perform functional enrichment analyses, identifying overrepresented Gene Ontology (GO) terms associated with domain combinations. This domain-level functional annotation provided a robust framework for understanding evolutionary adaptations and lineage-specific functional innovations.

### 7. Cellular localization and potential endosymbiotic origin of eukaryotic protein

To assess the potential endosymbiotic origin of selected eukaryotic proteins, we predicted their subcellular localization using TargetP2.0 [47]. This tool estimates the likelihood that a given protein is targeted to endosymbiotic organelles, specifically the mitochondria or chloroplasts, based on N-terminal signal sequences. TargetP2.0 was run with default parameters on the subset of eukaryotic proteins of interest. Predictions were used to infer whether domain combinations enriched in Eukaryota—and contributing to specific phylogenetic topologies—may be associated with organelles of endosymbiotic origin.

### 8. Ancestral Character Reconstruction

Ancestral states of domain-presence characters were inferred using a homemade R pipeline built around the ace() function in ape [48] and supplementary utilities in phytools [49]. Each binary trait—denoting the presence or absence of a specific Pfam domain or domain combination—was mapped onto a rooted species tree, and two Mk2 models were fitted: the equal-rates (ER) [50] and all-rates-different (ARD) variants. For every trait, the Akaike Information Criterion was used to choose the better-supported model, with an additional stability filter that defaulted back to ER whenever the ARD model exhibited implausibly high rate heterogeneity (e.g., rate ratios exceeding 100).

Once model selection was complete, the pipeline extracted the posterior probability of the “presence” state at each internal node and compiled these probabilities into a unified results table covering thousands of characters. Key nodes showing significant gains or losses of domain combinations were then highlighted directly on the phylogeny using iTOL [51] (Letunic & Bork 2021), providing an intuitive, interactive visualization of evolutionary transitions in domain combination content.

## Acknowledgement

This research was carried out as part of a PhD thesis within the µPlatypus Lab, whose support is gratefully acknowledged. DPD is funded by the Institut Pasteur de Lille, France; Le Conseil régional des Hauts-de-France (dispositif STaRS, convention 22008657); L’Université de Lille; and the France 2030 investment plan (convention R-ERCGEN-23-007-DEVOS). This work was also supported by the Moore–Simons Project on the Origin of the Eukaryotic Cell (Grant No. 9733, https://doi.org/10.37807/GBMF9733).

## Supporting information

**S1 Fig.** Scatter plots showing the number of protein domains (X axis) and the number of domain pairs (Y axis) for representative genomes across the three superkingdoms: Bacteria, Archaea, and Eukaryota. Each subplot corresponds to one superkingdom and data points are colored by phylum in Eukaryota and by supergroups in Bacteria and Archaea. In the Eukaryota panel, three distinct clusters are outlined: one comprising mainly unicellular eukaryotes and fungi, another including plants and simpler metazoans, and a third corresponding to higher metazoans.

**S2 Fig.** Collapsed ML-Tree done using IQ-TREE with binary substitution model (GTR2+F0+R10), ultrafast bootstrap 1,000 iterations and bnni option (UFBoot trees by nearest neighbor interchange (NNI)) using domain content matrices as input.

**S3 Fig.** Collapsed ML-Tree done using IQ-TREE with binary substitution model (GTR2+F0+R10), ultrafast bootstrap 1,000 iterations and bnni option (UFBoot trees by nearest neighbor interchange (NNI)) using combined pair and domain content matrices as input.

**S4 Fig.** Comparison of 2 topologies, on the left ML-Tree of Figure 1 and on the right, ML-Tree done using IQ-TREE with binary substitution model (GTR2+F0+R10), ultrafast bootstrap 1,000 iterations and bnni option (UFBoot trees by nearest neighbor interchange (NNI)) using combined pair and uni-domain content matrices as input, but without Bacteria.

